# The dung beetle microbiome complements host metabolism and nutrition

**DOI:** 10.1101/2025.05.02.651921

**Authors:** Joshua A. Jones, Armin P. Moczek, Irene L. G. Newton

## Abstract

Many multicellular organisms rely on communities of microbial organisms to properly benefit from their diets, for instance by assisting in the breakdown of complex polysaccharides, the synthesis of essential resources, detoxification, or even preventing putrefaction. Dung beetles commonly rely on herbivore dung as their main source of nutrition, a diet rich in recalcitrant, hard-to-digest plant polysaccharides yet poor in essential amino acids, which animals typically cannot synthesize on their own. The work presented here investigates the potential role of the host-associated microbial community in allowing these insects to thrive on their nutrient-poor diet. Specifically, we investigated if the microbiota of the bull-headed dung beetle, *Onthophagus taurus*, may be capable of synthesizing amino acids and breaking down complex plant polysaccharides. To do so we functionally annotated genes within metagenomically assembled genomes (MAGs) obtained via shotgun-metagenomic sequencing. The annotation of these MAGs revealed that bacteria found in association with *O. taurus* possess the metabolic potential necessary to bridge the gap between host metabolic needs and the limitations imposed by their diet. Specifically, *O. taurus* microbiota contain amino acid biosynthesis pathways and genes encoding cellulases and xylanases, both of which are absent in the beetle genome. Further, multiple functionally relevant bacterial taxa identified here have also been observed in other studies across diverse dung beetle species, possibly suggesting a conserved pool of dung beetle symbionts and metabolic functions.

**Importance:** Host-symbiont interactions allow animals to take advantage of incomplete and/or challenging diets and niches. The work presented here aims to identify the physiological and metabolic means by which host associated microbial species shape the ecology of one of the most speciose genera in the animal kingdom: dung beetles in the genus *Onthophagus*. Both larva and adult stages of most *Onthophagus* rely on herbivore dung, a diet rich in recalcitrant, hard-to-digest plant polysaccharides yet poor in essential amino acids, which animals typically cannot synthesize on their own. To utilize such a challenging diet, *Onthophagus* vertically transmit a maternally derived microbial community which supports normative development in immature individuals and maintenance and reproduction in adults. Taken together, *Onthophagus’* extraordinary diversity, complex ecology, and varied relationship with their microbial associates make them an ideal system to investigate mechanisms and diversification of host-diet-microbiome interactions.

## Introduction

Animals rely heavily on key symbionts within their microbiome to exist within their ecological niche. Depriving these animals of their microbes often renders them unable to use their focal food resource as their symbionts provide key metabolic pathways. For example, insects such as the rhinoceros beetles, termites, and leafcutter ants, lack the ability to reliably break down the complex plant polysaccharides in their diets^1–3^. Instead, these insects harbor bacterial or fungal symbionts with the metabolic potential to produce diverse enzymes to break down these resources and provide simpler components to the host. Conversely, many animals consume diets that are rich in simple carbohydrates yet poor in essential nutrients the animals cannot make. For example, aphids, bees, and stink bugs all consume diets rich in sucrose, fructose and glucose but poor in essential amino acids. However, in each case microbial symbionts are capable of synthesizing essential amino acids to the benefit of their host^4–6^. The importance of these interactions to animal resource use suggests that host-symbiont interactions are essential to understanding animal ecology and evolution.

Dung beetles, which specialize in feeding on the dung of other animals, are another clade of animals reliant on a challenging diet. Onthophagine beetles, specifically, are obligately reliant on herbivore dung, which presents further difficulty because of the abundance of tough to digest plant materials and relative lack of essential nutrients. Despite this, dung beetles are extraordinarily species-rich, with the genus *Onthophagus* alone accounting for an estimated 2,500 extant species^7^. Recent work suggests that onthophagine dung beetles may owe a portion of their evolutionary success to a community of heritable and functionally significant microbes. Inhibiting the inheritance of these microbes results in prolonged developmental time and decreased adult size^8^, suggesting that these beetles are reliant on their microbiome for normative development. This, in light of their difficult diet, has fueled the hypothesis that dung beetles rely on the metabolic pathways encoded within their microbial associates to efficiently utilize and complement their diets. To date, the only data supporting this hypothesis relies on either 16S rRNA based functional predictions^9,10^ which, however, often miss functional differences within taxa^11^, or the metabolic activity of isolated microbes^12^, which focuses on compounds that are likely digestible by host genomes^13^. Here, we set out to construct a metagenome of the symbionts of the bull-headed dung beetle *Onthophagus taurus* to test the hypothesis that the dung beetle microbiome encodes metabolic capabilities that empower dung beetles to utilize an otherwise hard-to-digest and incomplete diet.

To accomplish this, we sequenced the bacterial community from the *O. taurus* larval gut, the adult midgut, and the pedestal (a fecal pellet left by ovipositing mothers which assists in the passage of microbes across generations^8,14,15^). These reads were assembled into individual metagenomically assembled genomes (MAGs) and annotated to functionally and taxonomically characterize the dung beetle microbiome. Finally, the metabolic potential of the microbiome was compared across three onthophagine dung beetles (*O. taurus, Onthophagus sagittarius*, and *Digitonthophagus gazella*) to determine if functional deficits present within the host genomes are complemented by the genes present in the microbiome. Our results suggest that the dung beetle microbiome has the potential to compensate for deficiencies in the host genome with respect to both complex polysaccharide breakdown and essential amino acid synthesis. Further, taxonomic identifications provide evidence that Bacteroidales and Pseudomonadota may be particularly important in fulfilling these roles in *O. taurus*.

## Materials and Methods

### Sample preparation, sequencing, quality control, and assembly

Details on the specific methodologies used to produce samples, sequences, and MAGs are described in the companion Microbiology Resource Announcement. In brief, five libraries were produced by pooling samples across five sample types, larval for-, mid-, and hindguts, adult midguts, and pedestals. Importantly, these samples were enriched for bacteria (following methods modified from^16^) decreasing the representation of DNA from the host and other eukaryotes. A total of 32 MAGs were produced and 16 were determined to be >90% complete (using CheckM^17^) and uploaded to NCBI in BioProject PRJNA1117517 (see Table 1, for BioProject labels and SRA Accession numbers). The assemblies for the remaining 16 MAGs were uploaded to the DRYAD repository. Details on MAG assembly quality, content, identification, and the total proportion of assembly reads binning within the MAG are summarized in Table 1.

**Table 1:**
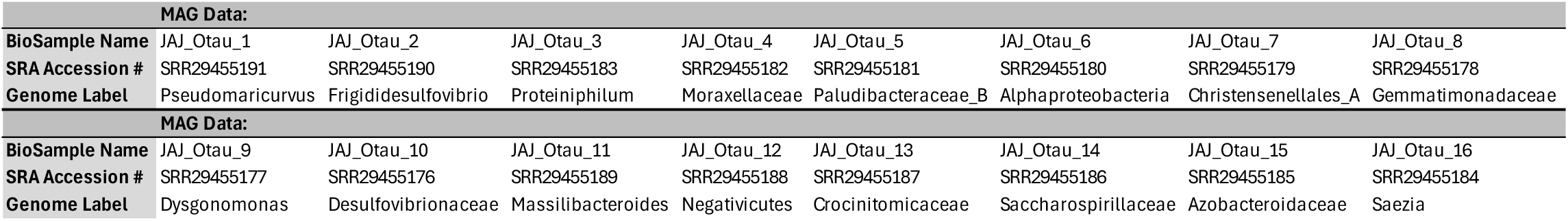
BioSample and SRA Accession numbers of genomes over 90% complete. Genome Label denotes how the sequences accessible through NCBI are denoted throughout this manuscript.

### Characterization of microbial communities

Community composition was determined for unassembled reads after initial quality control steps (removing host-associated DNA sequences and trimming low-quality reads). Reads were processed on KBase^18^ using Kaiju v1.9.0^19^, in greedy mode, and the Kaiju databases: RefSeq Complete Genomes (protein sequences from completely assembled bacterial, archaeal, and viral genomes from NCBI RefSeq updated March 23^rd^ 2022), and Fungi, protein sequences from a representative set of fungal genomes (updated March 23^rd^ 2022). Kaiju output contained the relative and total abundance of each taxon, no normalization or transformation was conducted. The resulting compositional data was transferred to R v4.2.2^20^ for figure production and to determine which taxa were shared across samples. The Kaiju outputs of each sample are deposited into the DRYAD repository.

### Determining amino acid synthesis pathway completion

Metabolic potentials for *Onthophagus taurus, Onthophagus sagittarius*, and *Digitonthophagus gazella* were all determined using DNA sequences and gene predictions from Davidson & Moczek 2024^21^. To avoid associating genes from any potential microbial contamination with the beetle genomes, analysis was limited to the definitive chromosomes. These include: chr1, chr2, chr3, chr5, Schr6, chr7, chr8, chr10, chr11, ScKx7SY_15, and ScKx7SY_16 for *O. taurus*, Scaffold_1, Scaffold_2, Scaffold_3, Scaffold_4, Scaffold_5, Scaffold_6, Scaffold_7, Scaffold_8, Scaffold_9, and Scaffold_10 for *O. sagittarius*, and ScIV947_1, ScIV947_2, ScIV947_3, ScIV947_5, ScIV947_6, ScIV947_7, ScIV947_9, ScIV947_12, ScIV947_14, and ScIV947_38 for *D. gazella*. The prediction of beetle amino acid synthesis pathways were determined by annotating the beetle genomes with the KEGG reference database^22^ and searching for amino acid synthesis genes in each genome. A pathway was considered complete if the enzyme catalyzing each reaction between pyruvate and that amino acid was present. The amino acid synthesis potential of the MAGs was determined with Gapseq^23^. Using the default parameters, Gapseq produced predictions for the completeness of amino acid biosynthesis modules. If a module was incomplete, any module requiring that one as a prerequisite was also considered incomplete.

### Annotating carbohydrate-active enzymes

The carbohydrate breakdown potential was quantified by using the KEGG annotations from the beetles and the RASTtk^24–26^ annotations from the MAGs. RASTtk annotations for all 32 MAGs are deposited into the DRYAD repository. Direct breakdown potential for major dung components (cellulose, cellobiose, xylan, and xylose) was quantified by counting related genes throughout these annotations. To determine further carbohydrate breakdown potential CAZymes (Carbohydrate-Active enZYmes) within the beetle genomes were identified using the dbCAN HMM database v12.0 and instructions found here (https://bcb.unl.edu/dbCAN2/download/Databases/dbCAN-old@UGA/)^27^. In short, HMMER v3.4^28^ (hmmer.org) was used to determine which genes within each beetle genome were annotated in the CAZyme database before a parser script was used to assign those genes into CAZyme families. In parallel, annotated MAGs were input into the dbCAN2 app^27,29,30^ (v1.9.1), on KBase, to produce similar annotations based on CAZyme families. We focused on the number of genes assigned to glycoside hydrolase (GH) families, quantifying their abundance across each beetle genome and MAG.

## Results

### Assembly results and statistics

Initial sequencing and quality control resulted in a total of 54,481,286 reads across our five libraries. These reads were assembled into a total of 26,749 contigs which binned into 32 MAGs. 28.4% of total reads mapped to these 32 MAGs or the host genome (20.9% and 7.53%, respectively), suggesting we captured only a subset of the total microbial diversity present in these environments within these MAGs. For this manuscript we focus on raw reads to estimate community composition, contigs to estimate overall metabolic potential, and the 16 nearly complete MAGs, with a completion score above 90% in CheckM^17^, alongside the 16 less complete MAGs, completion between 44.0%-89.8% for more specific functional predictions and annotations.

### Bacterial and fungal composition

To gain insights into any potential community differences across these sample types, we assessed the bacterial and fungal communities using unassembled reads characterized with Kaiju. The Greedy mode of Kaiju translates DNA into proteins and breaks reads into fragments before identifying a read based on the database sequence with the highest possible similarity score, allowing for substitutions^19^. This results in a higher rate of identification of genera underrepresented in the database and allows for novel genera to be identified to the family level if other genera are available^19^. Kaiju was able to classify 6-54% of reads as bacterial and 0.6-2.6% as fungal across samples, with the adult midgut having the lowest classification rate and the pedestal having the highest for both bacteria and fungi (Figure 1; Supplemental figure 1). The much lower classification rate of fungi was likely the result of our sample processing (i.e., enriching for bacterial reads to decrease host reads likely also decreased fungal reads) and not a representation of their differential abundance. Across all samples, Kaiju identified 1720 bacterial families, 65% of which were present in all samples (Supplemental figure 2). The fungal communities identified were less diverse overall, with 222 fungal families being identified, 85% of which were present in all samples. No single bacterial genus dominated any sample, but the most abundant genera were all Pseudomonadota or Bacteroidota (Figure 1).

**Figure 1:**
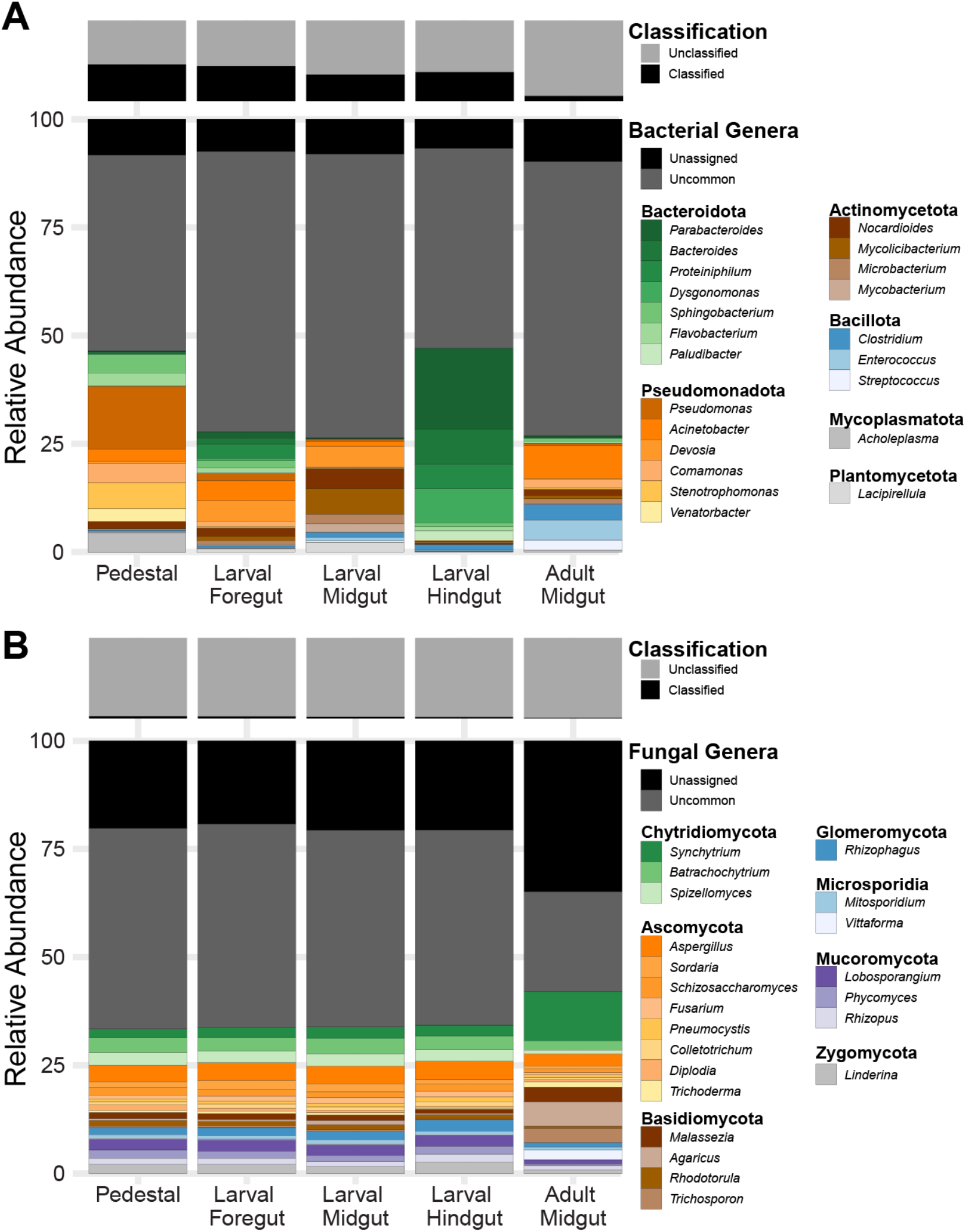
Relative abundance of bacteria and fungi across sample types. A) shows the relative abundance of the most common bacterial genera across samples and B) shows this data for fungi. “Fraction classified”, above relative abundances, shows the total percentage of quality-controlled reads from the sample that could be assigned to taxa in that group. Uncommon taxa were combined to form the <Others= category, and taxa that couldn9t be identiûed to the genera level were combined to form the <Unassigned= category.

### Deficits in host amino acid synthesis pathways are present in microbial genomes

As herbivore dung may be poor in essential amino acids, we determined complete amino acids synthesis pathways in the host and microbial genomes to ascertain if deficits in host metabolism could be supplemented through biosynthesis by microbes. All three dung beetle species (*O. taurus, O. sagittarius*, and *D. gazella*) appear to possess the same amino acid synthesis potential as other arthropods^31^, retaining the genes to encode enzymes necessary to synthesize only 10 of 20 amino acids (Figure 2a). The microbial genes represented throughout the MAGs have the potential to encode enzymes for synthesizing all 20 common amino acids. Saccharospirillaceae was the only MAG which contained all the genes necessary to synthesize all 20 amino acids, with the next highest potentials (18 amino acids) found in the Moraxellaceae and *Saezia* (17). The remaining MAGs varied in potential, encoding genes able to synthesize between two and sixteen amino acids. Thus, the onthophagine gut microbiome appears to contain multiple copies of amino acid synthesis pathways not found in host genomes.

**Figure 2:**
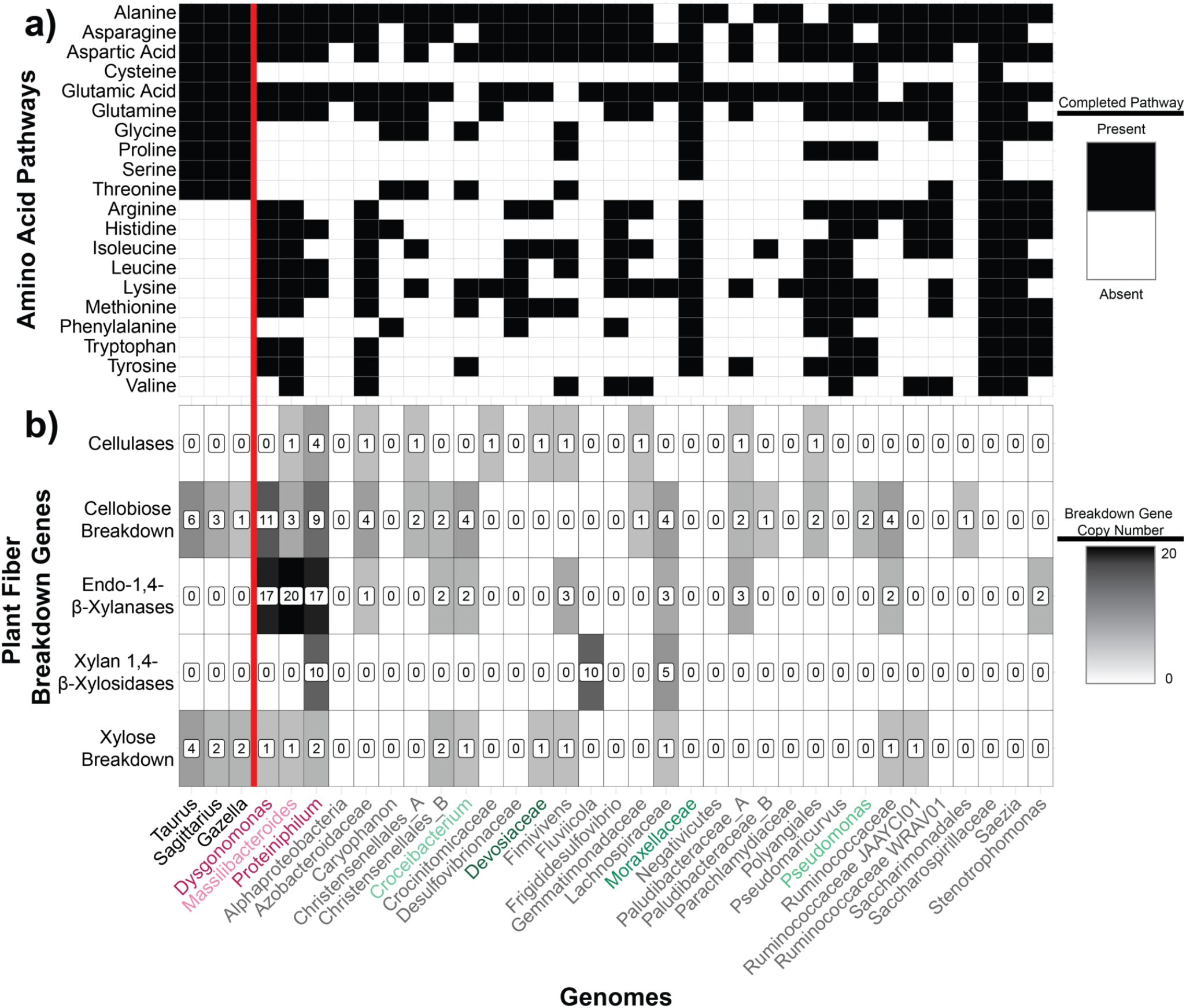
Metabolic potentials of beetles and gut bacteria. a) Presence/absence of completed amino acid pathways. Each beetle genome contains the genes to synthesize 10 of the 20 essential amino acids while metabolic modelling predicts that microbes present in *O. taurus* can synthesize all 20. b) Abundance of putative cellulose and xylan breakdown genes. Bacteria found in *O. taurus* contain genes to breakdown cellulose, xylan, and their simpler components while beetle genomes only contain the genes to breakdown the simple components. Bold genome names represent beetle genomes, colored genomes names are in families represented in Figure 1, and grey names are those from families that were in the <Other= group in Figure 1.

**Figure 3:**
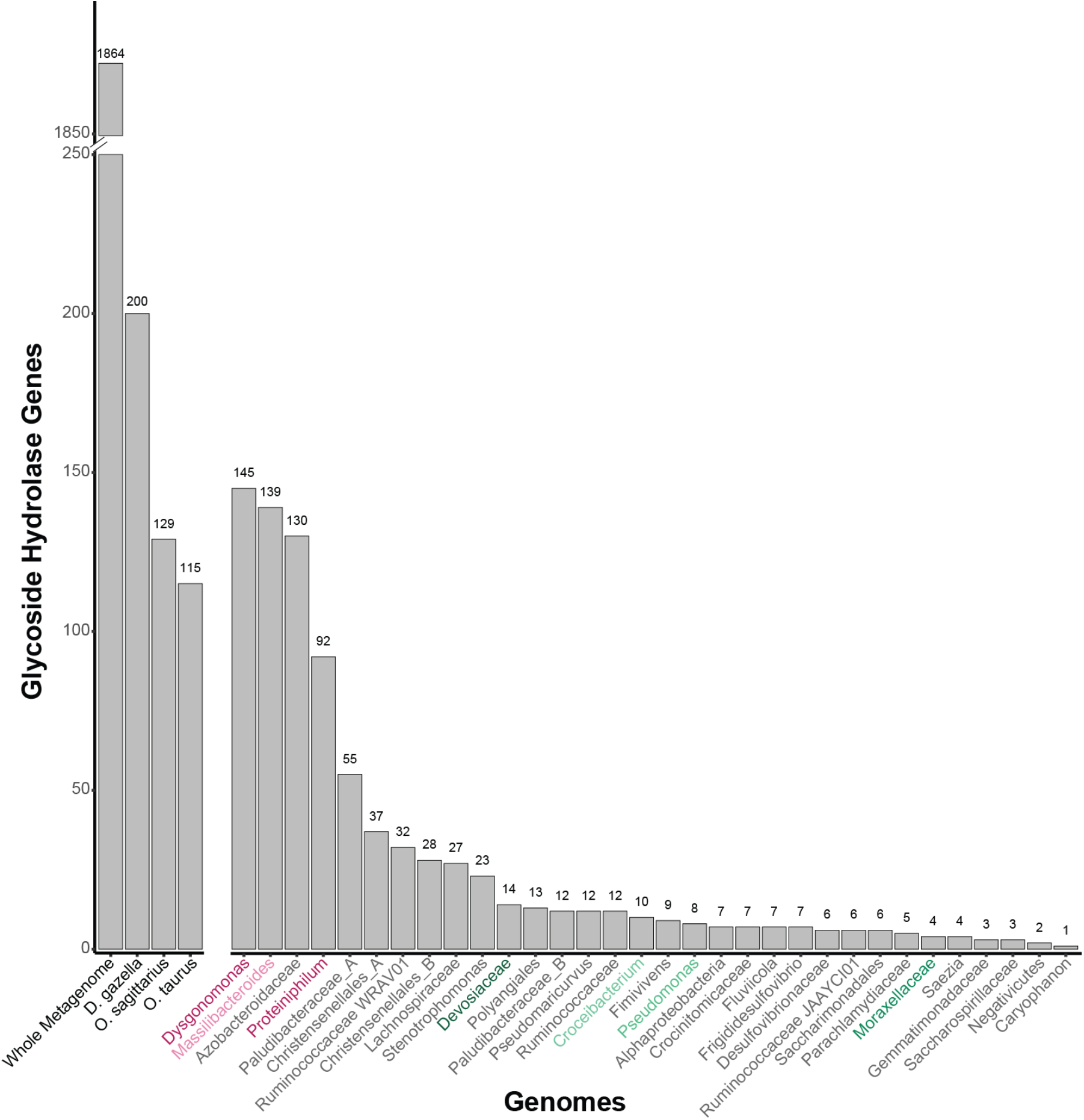
Abundance of putative glycoside hydrolase genes within the whole metagenome, individual beetle genomes, and individual MAGs. *D. gazella* contains far potential GH genes than *O. sagittarius* or *O. taurus* as well as the individuals MAGs, which have abundances similar to *O. sagittarius* and *O. taurus*. The counts within the beetle genomes, as well as the individual MAGs, are all dwarfed in comparison to the culmination of potential GH genes throughout the entire metagenome.

### Bacterial genomes contain complex polysaccharide breakdown genes absent from host genomes

The dung diet comprises complex polysaccharides from plant cell wells, often recalcitrant to animal digestion. To understand how the beetle may overcome this obstacle we identified genes encoding enzymes which may breakdown these complex polysaccharides or their simpler components. Neither the *O. taurus, O. sagittarius*, nor *D. gazella* genomes contained genes homologous to any cellulase nor xylan-breakdown genes (Figure 2b). Yet all three genomes contain homologous genes to enzymes which break down the simpler components of those polysaccharides, cellobiose (n = 6, 3, and 1, respectively) and xylan (n = 4, 2, and 2, respectively). In contrast, dbCAN2 identified 13 putative cellulases and 97 putative xylan-breakdown genes across the 32 MAGs. The majority of the xylan-breakdown genes (n = 72) were endo-1,4-beta-xylanases with the remaining (n = 25) being xylan 1,4-beta-xylosidases. Although the MAGs encoded more enzymes associated with xylan breakdown than cellulose breakdown, we identified more genes associated with the degradation of cellobiose (n = 52) than xylose (n = 12). Across all MAGs, *Proteiniphilum* encoded the largest number of genes across these categories with 31 total cellulose or xylan break-down genes, followed by *Massilibacteroides* with 21, and *Dysgonomonas* with 17.

To get an idea of the broader polysaccharide breakdown potential, we also quantified the total number of predicted glycoside hydrolase (GH) genes across both host genomes and MAGs. GHs are a broad group of enzymes defined by their potential to hydrolyze glycosidic bonds. This family of genes includes those capable of breaking down cellulose, cellobiose, xylan, and xylose, as discussed above, in addition to many other carbon sources. By quantifying GHs we were able to achieve an estimation of the potential for these organisms to digest a broader variety of complex polysaccharides which they may encounter. We found that *D. gazella* possesses the highest GH count (200), whereas *O. sagittarius* and *O. taurus* featured more similar counts at 129 and 115, respectively. However, these values are dwarfed by the GH abundance across all assembled contigs (n=1,864) and are comparable to the highest values within individual MAGs (*Dysgonomonas* = 145, *Massilibacteroides* = 139, and *Azobacteroidaceae* = 130). More generally, these results suggest that the microbial community may expand the ability for the dung beetles to derive nutrition from their diet by encoding an abundance of polysaccharide-utilization genes.

## Discussion

Many animals that consume complex or nutrient poor diets rely on microbial symbionts for digestion or nutrient synthesis. Onthophagini beetles obligately consume herbivore dung as a diet throughout development and into adulthood^14^. Herbivore dung is a challenging diet because it is mostly composed of tough plant materials which animals often can’t digest^1,2,32^. Additionally, essential amino acids, which animals must consume from their diet, are also rare in herbivore dung^32^. Together, this makes dung a particularly difficult food source and suggests that dung beetles may require their microbes to both digest and synthesize essential resources from this diet. Here, we combined analysis of the metabolic potential of three onthophagini beetle genomes (*O. taurus, O. sagittarius, D. gazella*) with that derived from shotgun-metagenomic sequencing of *O. taurus* gut sections and pedestals. By comparing these datasets, we validated expectations of metabolic deficits within the host genomes while simultaneously identifying candidate microbial symbionts to supplement these deficits.

### Evidence for diverging microbial communities across gut samples

Characterizations of the bacterial and fungal communities derived from different gut sections and host developmental stages revealed several interesting patterns. Notably, the fungal community structure was more stable between sample types than that of the bacterial community, with the majority of fungal species found across all samples. This result, obtained in this study on *Onthophagus* beetles, contrasts with previous work in another dung beetle species showing strong differentiation in fungal communities in the gut across the life stages of the dung beetle *Catharsius molossus*^33^, suggesting possible differences in host-fungal interactions across the species. Compared to fungal abundance, the relative abundance of bacterial taxa across samples was more dynamic. For example, Dysgonomonadaceae and Moraxellaceae, bacteria which have previously been shown to be abundant in numerous dung beetle genera^9,10,15,33–35^, exhibited pronounced differences in relative abundance depending on sample. Specifically, Dysgonomonadaceae was found in relatively robust abundance in the larval fore- and hindgut (4.59% and 15.47%) but was rare in the pedestal and larval and adult midguts (0.33%, 0.07%, 0.41%) while Moraxellaceae was present in the pedestal, larval foregut, and adult midgut (3.00%, 4.67%, 7.62%), but comparatively rare in the larval midgut and hindgut (1.06%, 0.11%). The presence of *Dysgonomonas* across different dung beetle species may be indicative of a shared and conserved function in digestion. Further, the abundance of *Dysgonomonadaceae*, a wide-spread and potentially beneficial family, in the hindgut fits with previous research in *Pachnoda* which suggests that scarab beetles may have the highest densities of beneficial microbes in the hindgut^13,37,38^. All together, these results suggest that dung beetles likely harbor the highest densities of their beneficial symbionts within the hindgut, akin to many other insects reliant on their symbionts for digestion^1,39–42^, and that *Dysgonomonas* specifically may be a functionally relevant hindgut inhabitant. Finally, the differences we observe across samples, and previous observations of strong differences across life stages^9,15,33^, suggest that host-derived factors likely play a major role in microbial community assembly. Future work may seek to confirm these gut section dynamics with higher replication and methods allowing for increased fungal representation.

### Evidence for reliance on microbiome members for resource digestion and synthesis

We sought to determine if onthophagini dung beetles use their symbiotic microbes for the digestion of complex polysaccharides abundant in their diet. Our results show that, like many animals, the beetles lack the genes necessary for the production of enzymes to break down either cellulose or xylan, two polysaccharides abundant in herbivore dung^43^. Despite this, the beetle genomes do contain genes related to the breakdown of the cellobiose and xylose, simpler components of those complex polysaccharides. Together this indicates a gap in the metabolic potential of *Onthophagus* dung beetles to digest their diet. This gap, however, may be filled by the gut microbial community, which encodes an abundance of enzymes able to break down these complex polysaccharides. The majority of these genes are related to xylan breakdown, suggesting that the bacteria may specialize on the hemicellulose within the dung, and not the cellulose itself. That said, the microbiome does also encode cellulose breakdown enzymes, including an abundance of diverse glycoside hydrolases, suggesting broader carbohydrate breakdown potential beyond this specialization. Interestingly, many of these xylan, cellulose, and carbohydrate breakdown genes were present within Bacteroidales, namely *Dysgonomonas, Massilibacteroides, Proteiniphilum*, and *Azobacteroidaceae. Dysgonomonas*, as mentioned above, appear to be prevalent across dung beetles, having been observed in *Copris incertus, Catharsius molossus, Euoniticellus intermedius* and *E. triangulatus, E. fulvus, O. binodis, O. australis, O. hecate, E. fulvus*, and multiple populations of *O. taurus*, as well as within each *O. taurus* life stage^9,10,15,33–36,44^. The consistent occurrence of *Dysgonomonas* across dung beetles, along with the functional potential described here and in other systems^45–49^, suggests that *Dysgonomonas* may be particularly important to dung beetle ecology. Overall, these results support the prediction dung beetles likely rely on their microbiome to digest the complex polysaccharides abundant in their diet and highlights some bacteria with that may be key to the maintenance of this function.

Further, our results suggest that onthophagini beetles may also rely on a functionally redundant set of symbionts to synthesize amino acids from their diet. Analysis of the beetle genomes confirmed that they lack the synthesis pathways to produce 10 of the 20 amino acids, while corresponding synthesis pathways for all 20 of these amino acids are represented within multiple MAGs. The symbionts with the highest number of complete pathways belonged to Pseudomonadota, some of which encoded enzymes able to synthesize the majority (Moraxellaceae and *Saezia*), or all (Saccharospirillaceae), of the essential amino acids. Interestingly, several Bacteroidales MAGs (*Dysgonomonas, Massilibacteroidales*, and *Azobacteroidaceae*) also contained the majority of the synthesis pathways missing from the host genomes (8, 9, and 9, respectively). Yet, MAGs assembled from Bacteroidales contained fewer total complete pathways than those assembled from Pseudomonadota, with *Dysgonomonas* containing 13, *Massilibacteroidales* containing 14, and *Azobacteroidaceae* containing 14 compared to the 20, 18, and 17 of Saccharospirillaceae, Moraxellaceae, and *Saezia*, respectively. The potential role of Pseudomonadota in dung beetle ecology may thus be important more broadly, as Moraxellaceae, in particular *Acinetobacter*, has been shown to co-occur with *Dysgonomonas* across many dung beetle species and life stages^9,10,15,33–35,44^. Interestingly, a survey of the *O. taurus* microbiome suggests that Pseudomonadota may be highly abundant within the dung yet rare in the developing larvae^15^, raising the possibility that Pseudomonadota may be a reliable provider of amino acids, independent of their potential to colonize within or be horizontally inherited by the host.

### Conclusion

We investigated the potential role of the microbiome in complementing the metabolic potential of dung beetles in the face of a challenging and incomplete diet. Specifically, we find that the three beetle genomes surveyed lack genes encoding enzymes necessary for breaking down complex polysaccharides or synthesizing 10 of 20 amino acids, key metabolic deficiencies given their cellulose- and xylan-rich yet amino acid-poor diet. However, the genes harbored within the gut microbiome encode the complementary metabolic potential to bridge this gap and transform dung into a resource beetles can use. Furthermore, we identify specific bacterial taxa, some of which have been found across multiple other studies in diverse dung beetle species, which may be especially important in fulfilling this function. Together this work demonstrates how the microbiome may have scaffolded the origin and diversification of dung beetles into their unique niche. Future studies are needed to now assess the precise contributions of select microbiome members *in vivo*, and if and how microbiome diversity interacts with host diversity in development and evolution.

## Data Availability

The project discussed here has been deposited in NCBI under BioProject PRJNA1117517. Included in this are the raw sequence reads from the pedestals, larval fore-, mid-, and hindguts, and the female adult midguts (accession no.: SRR29215567, SRR29215566, SRR29215565, SRR29215564, & SRR29215563, respectively). Further, this project also includes the reads for all 16 MAGs discussed above (accession no.: SRR29455191, SRR29455190, SRR29455183, SRR29455182, SRR29455181, SRR29455180, SRR29455179, SRR29455178, SRR29455177, SRR29455176, SRR29455189, SRR29455188, SRR29455187, SRR29455186, SRR29455185, & SRR29455184). Reads for the 16 lower-quality MAGs, summary statistics for all MAGs, and the protein annotations for all MAGs are included in the DRYAD repository (https://doi.org/10.5061/dryad.4qrfj6qn8).

## Acknowledgements

This work was conceived and written by JAJ, IGN, and APM with JAJ and IGN carrying out experiments and JAJ conducting data processing and analysis. We would like to thank Tracy Rettig for collecting *O. taurus* in the field, Eve Piery, Isabel Manley, and Sadie Kidd for expert help with beetle husbandry, and the members of the Moczek and Newton Labs for constructive comments. This work was supported in part through generous funding from the National Science Foundation [Grant no. 2243725 and 1901680 to APM] and a National Science Foundation Graduate Research Fellowship [Grant No. 2141416 to JAJ]. The opinions, interpretations, conclusions, and recommendations are ours and are not necessarily endorsed by the National Science Foundation.

## Figures

**Supplemental Figure 1:**
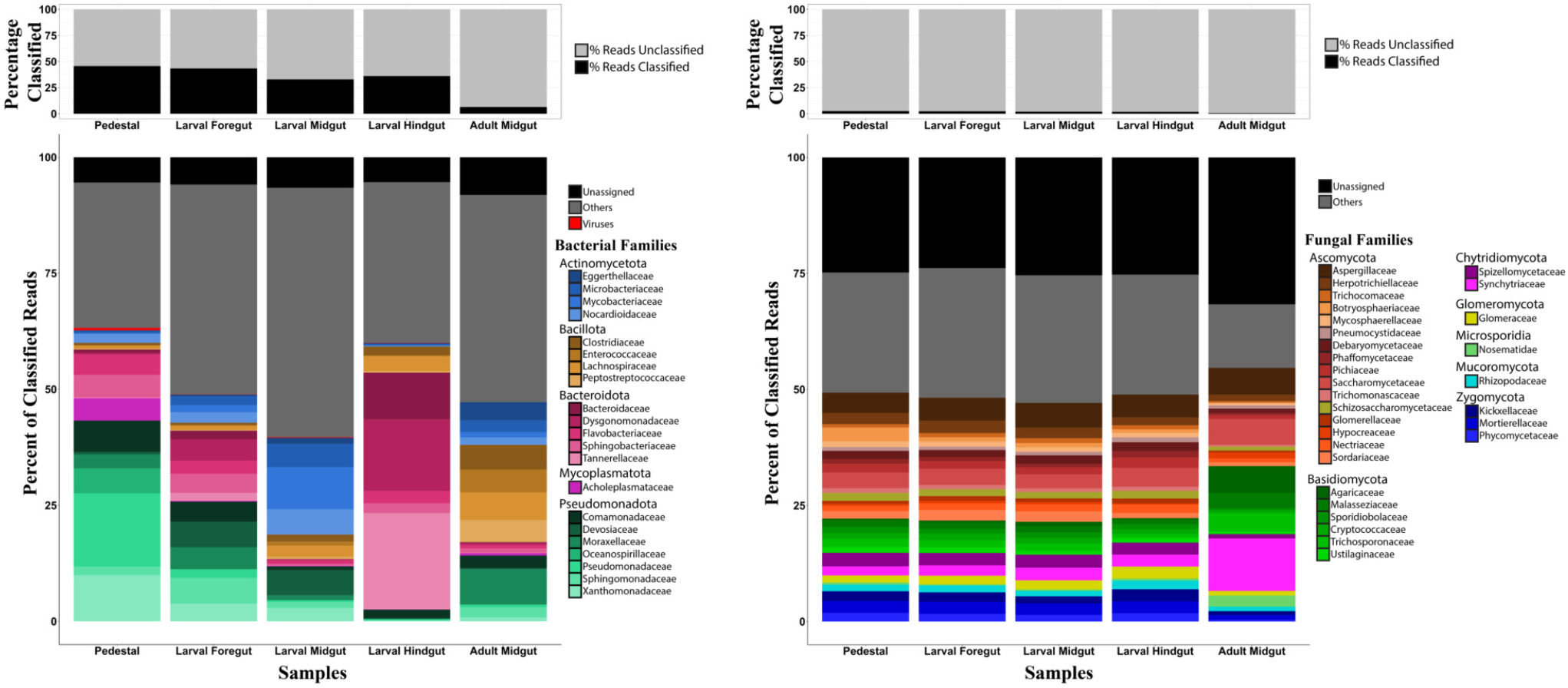
Relative abundance of bacteria and fungi across sample types. Graphs on the left show data for bacteria and graphs on the right show data for fungi. “Fraction classified”, at the top, shows the total percentage of quality-controlled reads from the sample that could be assigned to taxa in that group. Relative abundance graphs show families representing over 10% relative abundance of any sample, with the remaining classified families combining to form the “Others” category, and all taxa that couldn9t be identified to the family level being combined to form the “Unassigned” category.

**Supplemental Figure 2:**
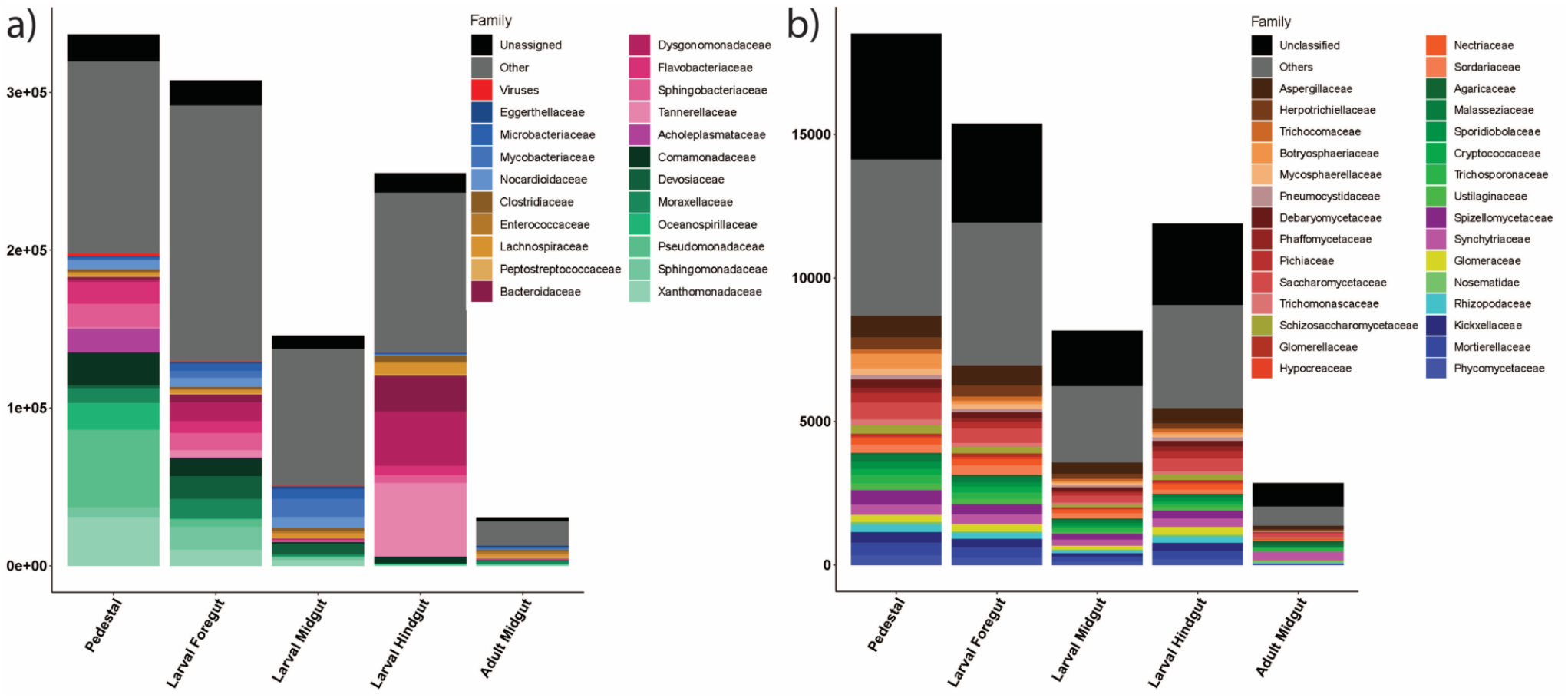
Absolute abundance of a) bacterial and b) fungal families across samples.

**Supplemental Figure 3:**
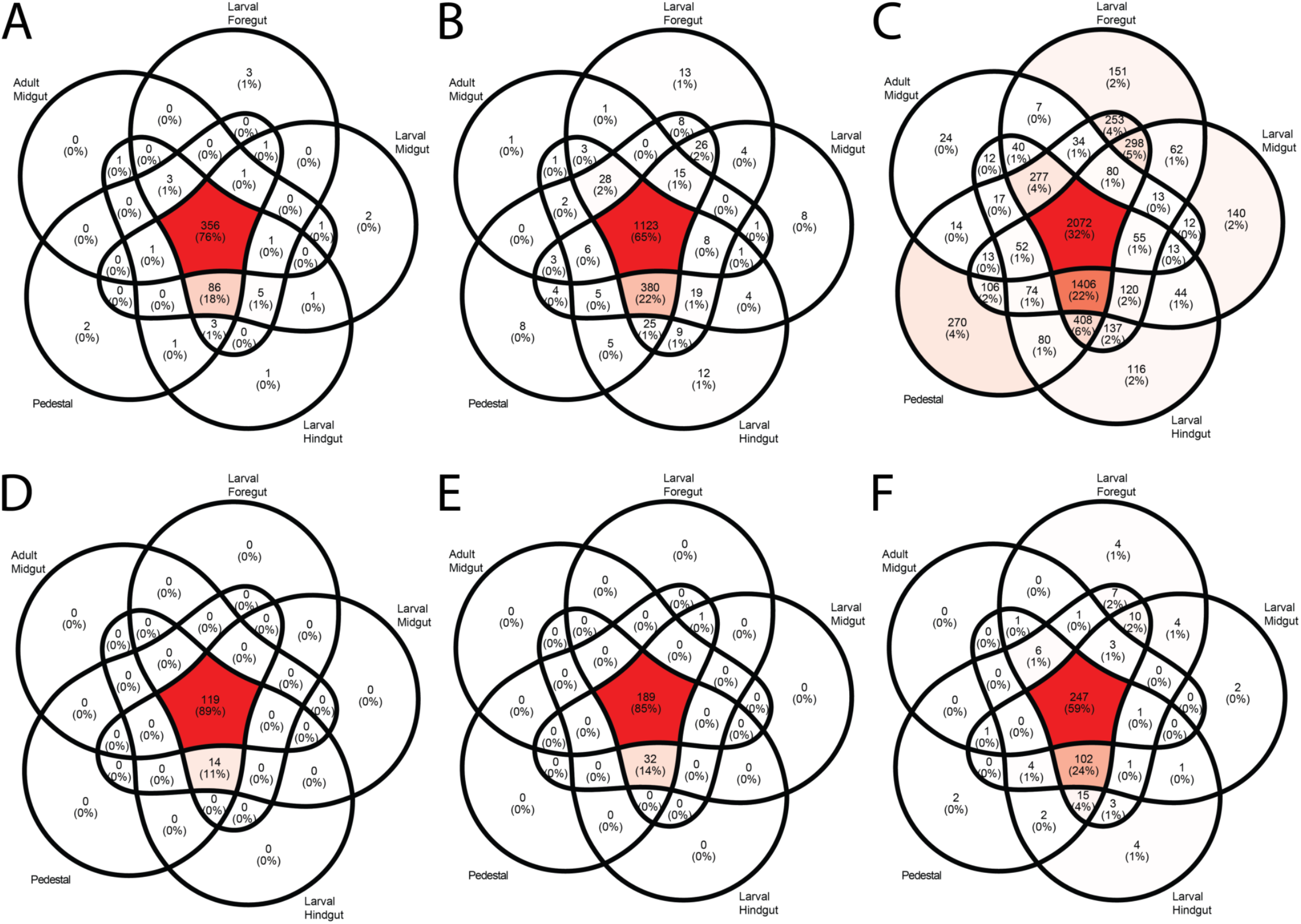
Bacterial and fungal taxa shared across samples. A-C are bacterial families, genera, and species, respectively, and D-F are fungal families, genera, and species. The majority of taxa are found across all samples, with the exception of bacterial species. In all cases the next highest percentage of taxa are found in all samples except the female adult midgut.

